# Exact formula of the total quasi-steady state approximation in competitive target-mediated drug disposition

**DOI:** 10.64898/2026.01.10.698771

**Authors:** Taehong Kim, Jaeyun Cha, Hyeseon Jeon, Hwi-yeol Yun, Dongju Lim, Jae Kyoung Kim

**Author notes:** These authors contributed equally (Taehong Kim, Jaeyun Cha, Hyeseon Jeon). Correspondence: Hwi-yeol Yun, PhD, College of Pharmacy, Chungnam National University, Daejeon, Republic of Korea, Dongju Lim, Department of Mathematical Sciences, KAIST, 291 Daehak-ro, Yuseong-gu, Daejeon, 34141, Korea, Jae Kyoung Kim, PhD, Department of Mathematical Sciences, KAIST, 291 Daehak-ro, Yuseong-gu, Daejeon, 34141, Korea.

## Abstract

Competitive target-mediated drug disposition (competitive TMDD) arises when two drugs compete for the same target receptor. These dynamics can be characterized by the full competitive TMDD model; yet, its complexity motivated the use of a reduced model, which is invalid under high receptor concentrations. While this problem can be resolved by using the total quasi-steady state approximation (tQSSA), which remains valid for all receptor conditions, the exact formula of the tQSSA-based reduced model—competitive qTMDD—has remained unknown for 15 years, as it requires solving a cubic equation and the nontrivial task of identifying a biologically meaningful solution. Consequently, researchers have relied on numerical approximation methods, which impose substantial computational costs to maintain accuracy and thereby hinder their practical use in complex real-world applications. To address this problem, we derive—for the first time—a real-valued exact formula for competitive qTMDD by leveraging analytic properties of cubic equations and geometric characteristics of their roots in the complex plane. This exact formula improved computational speed by more than 11-fold compared to previous numerical approximation methods, thereby enabling Bayesian inference using competitive qTMDD, which had been impractical due to excessive computational time. When applied to real-world data from clinical trials, competitive qTMDD estimated pharmacological parameter estimates comparable to those from the full competitive TMDD model while requiring only 30-43% of the computation time. Importantly, this estimation using competitive qTMDD remained consistently accurate regardless of data sparsity, whereas the previous reduced model produced biased estimates under sparse sampling conditions. By ensuring precise biological interpretation of drug systems even in complex real-world scenarios, the exact formula of competitive qTMDD has the potential to significantly streamline the drug development and clinical testing process. Our exact formula also consists entirely of real-valued terms, allowing seamless integration into existing pharmacometrics software.

**Author Summary:** Competitive target-mediated drug disposition (competitive TMDD) occurs when two drugs compete for the same target receptor. Analyzing this interaction has faced a dilemma for 15 years: choosing between a ‘full model’ that is accurate but computationally intensive, and a ‘reduced model’ that is fast but often loses accuracy under real-world clinical conditions. Researchers tried to solve this dilemma by deriving an accurate reduced model; however, it was mathematically challenging. As a result, they had to rely on numerical approximations—which require substantial computational power to maintain the accuracy needed for clinical use, making them impractical in real-world scenarios. In this study, we derive a first-ever exact formula for the new reduced model that is accurate across all biological conditions. This formula computes more than 11 times faster than previous numerical methods, making advanced statistical analyses—such as Bayesian inference—feasible for the first time in this context. When applied to real-world clinical data for Anakinra and rhIL-7-hyFc, our method yielded parameter estimates as accurate as the full model but required significantly less computation time. This breakthrough provides a more accurate biological interpretation and better guidance for determining the right drug dose, potentially accelerating drug development and reducing associated costs.

## Introduction

Competitive target-mediated drug disposition (competitive TMDD) describes the process where two drugs competitively bind to the same receptor (1–5). For example, when administering Immunoglobin G (IgG) for therapeutic purposes to patients with primary immunodeficiency (PID), endogenous IgG and therapeutic IgG competitively bind to a single receptor, the neonatal Fc receptor (FcRn) (1). Ignoring this competition introduces bias into predictions of drug efficacy (6). In particular, excluding endogenous ligands from the model underestimates the impact of drug disposition on the concentration of the ligand-receptor complex and its downstream reactions, hindering accurate efficacy prediction. Therefore, applying a competitive TMDD model is crucial to accurately reflect physiological characteristics.

The competitive TMDD model is represented by a system of ordinary differential equations (ODEs) that describes interactions between two competing drugs, a receptor, and their complexes. This system is often simplified using the quasi-steady state approximation (QSSA) to reduce the number of variables, thereby increasing computational efficiency and enabling parameter estimation with limited data (7). Previously, standard QSSA (sQSSA) was widely used (8–12); however, it fails to provide an accurate approximation when the total receptor concentration is higher than the drug concentration (12–15)—a situation frequently encountered in cases such as the interaction between Omalizumab and Immunoglobulin E (16). To address this limitation, the total QSSA (tQSSA) has been proposed as an alternative that achieves greater accuracy across a broader range of conditions (7, 17–22).

Despite its high accuracy, applying tQSSA to a competitive TMDD system is challenging, as it requires solving a cubic equation and selecting the biologically meaningful solution (i.e., the solution where all variables have positive concentrations) among the three possible roots (1, 20, 23). Although the smallest root is proven to be biologically meaningful, identifying the smallest root remains nontrivial (1). Consequently, the tQSSA formulation of competitive TMDD—competitive qTMDD—has relied on numerical approximation methods since its initial development nearly 15 years ago (1–3). However, these methods impose substantial computational costs to maintain accuracy, limiting their practical use in real-world applications. This underscores the urgent need for deriving an exact formula of competitive qTMDD to achieve both accuracy and efficiency in real-world competitive qTMDD applications

In this study, we derived an exact formula for competitive qTMDD by exploiting the geometric properties of the cubic roots in the complex plane. This formula enables an accurate approximation of the competitive TMDD system while achieving computation speeds 10-fold faster than previous numerical methods. As a result, parameter estimation, which previously took over 5.5 hours, could be completed in just 15 minutes of computation. Furthermore, when we applied competitive qTMDD to real-world Anakinra and rhIL-7-hyFc data, we accurately estimated the pharmacological parameters. In particular, the performance of competitive qTMDD remained consistently accurate regardless of data sparsity, contrasting the substantial inaccuracy observed in sQSSA-based simplification under the sparse data conditions. By providing a highly efficient and accurate means of parameter estimation for competitive drug systems, the exact formula of competitive qTMDD has the potential to enhancing the accuracy of drug dosing and biological interpretation of drug systems, consequently reducing the time and costs associated with drug development and testing.

## Results

### The exact formula for competitive qTMDD is essential

Target-mediated drug disposition (TMDD), the process by which a drug binds to its target site (or receptor) (24), is typically modeled using a nonlinear system of ordinary differential equations (Fig. 1(a), see Methods for details) (25–30). This model has been used in a simplified form to enhance computational efficiency and reduce model parameters. Among various simplification methods, one of the most widely used methods is the quasi-steady state approximation (QSSA) (Fig. 1a) (7). Applying standard QSSA (sQSSA) to the TMDD model yields the Michaelis-Menten TMDD model (mTMDD) by solving a simple linear equation for the drug-receptor complex (*RC*) (Fig. 1a (i)). In contrast, the quasi-steady state TMDD model (qTMDD), which is simplified with the total QSSA (tQSSA), involves solving a quadratic equation for RC and selecting the biologically meaningful solution (i.e.,0 < *RC* < *min*(*T, R*_*tot*_), where *T* is the total drug concentration and *R*_*tot*_ is the total receptor concentration among the two roots (Fig. 1a (ii)) (see Methods for details). Both mTMDD and qTMDD accurately approximate the TMDD when *R*_*tot*_ is low (Fig. 1b, left). In contrast, if *R*_*tot*_ is high (Fig. 1b, left), the accuracy of mTMDD deteriorates, while qTMDD remains accurate. In summary, for single-drug TMDD, the exact formula of both mTMDD and qTMDD can be derived, with qTMDD offering higher accuracy.

**Figure 1.**
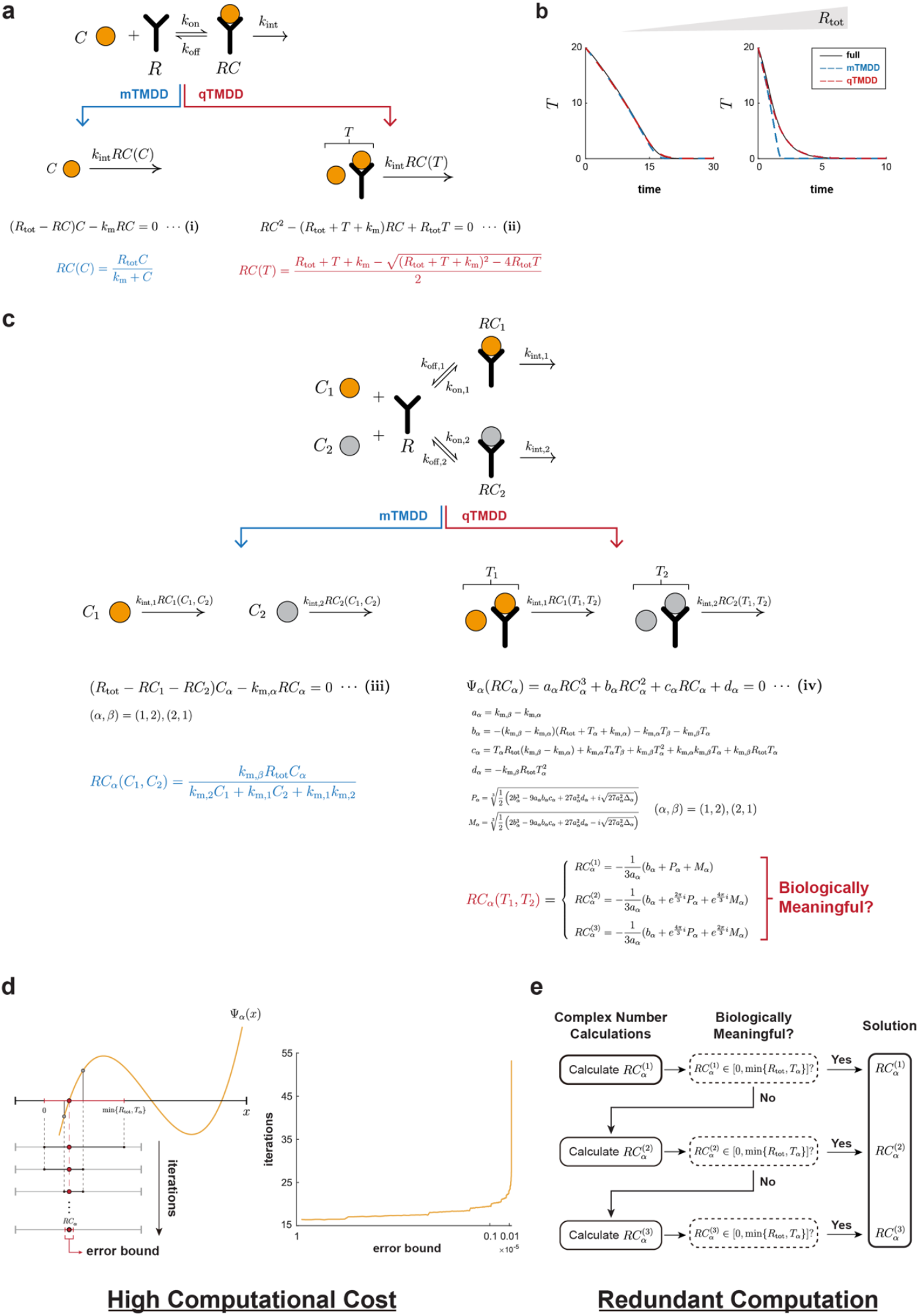
The exact formula of competitive qTMDD is largely unknown, leading to the use of inefficient alternative methods. **(a)** Single drug TMDD describes the binding of the drug (C) with the receptor (R), forming the drug-receptor complex (*RC*). This system can be simplified into two models: mTMDD and qTMDD. mTMDD only requires solving a simple linear equation (i), whereas qTMDD involves solving a quadratic equation (ii). This quadratic equation leads to two candidate solutions (*RC*^(1)^ and *RC*^(2)^), among which the unique biologically meaningful solution (i.e. the positive solution smaller than the total drug concentration (*T* = *C* + *RC*) and total receptor concentration (*R*_tot_ = *R* + *RC*) can be easily determined. **(b)** The plot showing the trajectory of total drug concentration (*T* = *C* + *RC*) simulated from the full TMDD (black line), mTMDD (blue dashed line), and qTMDD (red dashed line) under the low total receptor concentration (left) and high total receptor concentration (right). While the simpler mTMDD is accurate under the low receptor concentration, it becomes inaccurate at high total receptor concentrations. In contrast, the qTMDD remains accurate under both conditions. **(c)** Competitive TMDD describes the binding of two drugs (*C*_1_, *C*_2_) with the receptor (R), competing with each other to form the drug-receptor complex (*RC*_1_, *RC*_2_). This system can also be simplified into mTMDD and qTMDD models. mTMDD only requires solving a linear system of equations (iii), whereas the qTMDD involves solving two cubic equations (iv). Each of the cubic equations leads to three candidate solutions (*RC*^(1)^, *RC*^(2)^and *RC*^(3)^), but unlike the quadratic equation case (ii), distinguishing the unique biologically meaningful solution is mathematically unresolved, making an explicit form of competitive qTMDD remain unknown. **(d)** Consequently, the solution for the competitive qTMDD has been calculated relying on alternative methods. One representative example is the bisection method, which iteratively approximates the solution until reaching a desired error bound (left). However, as the error bound becomes stricter, the computational time increases exponentially (right). **(e)** Another method, the comparison method, calculates each of three candidates and identifies the biologically meaningful one. However, this method requires complex number computations, which can be challenging to implement in pharmacokinetic software such as NONMEM. In addition, the comparison method involves redundant calculations, increasing computational time.

The competitive TMDD model, which describes the competition of two drugs that bind to an identical target, can also be simplified using QSSA (Fig. 1c). The competitive mTMDD simplified with sQSSA can be derived by solving a system of linear equations for the two drug-receptor complexes (*RC*_1_, *RC*_2_) (Fig. 1c (iii)). However, this competitive mTMDD exhibits inaccurate approximation when total receptor concentrations are high (i.e., *C*_0_ + *K*_*m*_ ≪ *R*_*tot*_; Fig. S1), similar to the single-drug case. Given that single drug TMDD was more accurately approximated with tQSSA, its application to the competitive TMDD model, namely competitive qTMDD, is essential. Deriving the competitive qTMDD requires solving a cubic equation for *RC*_1_ and *RC*_2_ (Fig. 1c (iv)). This cubic equation yields three roots; however, identifying the biologically meaningful solution among them is nontrivial due to the presence of complex terms (Fig. 1c (v)). To address this, ae previous study (1) proved that the smallest of the three real roots is the unique biologically meaningful solution, but did not provide a method to identify the smallest root. Consequently, the exact formula of competitive qTMDD remains unknown even while it provides a more accurate approximation than competitive mTMDD (S1 Fig).

As the exact formula of competitive qTMDD is unknown, its simulation has relied on numerical approximation schemes solving cubic equations (Fig. 1d-e). For instance, the bisection method (Fig. 1d), which finds the approximate root in a specific interval, has been used (see Methods for more details). Starting from the interval where the biologically meaningful solution is guaranteed to exist (i.e., 0 < *RC* < *min*(*T, R*_*tot*_)), the solution is approximated by bisecting the intervals into halves (Fig. 1d, left). However, reducing the error of this approximation as much as desired requires numerous iterations, leading to long computation times. Indeed, the computation time (i.e., number of iterations needed to approximate the solution within the error bound) increases exponentially as the error bound decreases (Fig. 1d, right). Therefore, using the bisection method to compute competitive qTMDD is highly inefficient.

As an alternative, the comparison method (Fig. 1e), which calculates all three roots of the cubic equation using the cubic formula involving complex numbers, and checks whether each solution is biologically meaningful, has also been used (see Methods for more details). However, the major limitation of this method is its reliance on complex number arithmetic, which is not compatible with real valued data by the non-linear mixed effect modeling (NONMEM) method, widely implemented in field of pharmacometrics. While this issue can be solved by using another numerical approximation algorithm (see Methods for more details), this algorithm requires massive computation. As a result, the comparison method is also impractical, highlighting the urgent need for an exact formula of competitive qTMDD.

### We derived the exact formula of competitive qTMDD

To derive the exact formula of competitive qTMDD, (1) showed that the biologically meaningful solution is the smallest root among the three roots of the cubic equation. (See Lemma 1-4 in S1 Text). However, identifying the smallest root analytically proved challenging. As a result, despite the theoretical insights, the computation of competitive qTMDD continued to rely on numerical approaches such as the bisection or comparison methods.

To overcome this limitation, we found the exact formula of competitive qTMDD by explicitly identifying the smallest root among the three roots (Fig. 2a; see Methods for more details). We first derived the exact form of the concentration of drug-receptor complex associated with the stronger binding affinity (Fig. 2b (i); see Theorem 1 for more details). For instance, if drug *C*_1_ has a stronger binding affinity than drug *C*_2_, then the smallest root of the cubic equation for *RC*_1_ can be analytically obtained (Fig. 2b (i)). Then, this was reformulated to contain only real-valued terms (Fig. 2b (ii); see Theorem 1). This formula of *RC* with stronger binding affinity was substituted into the *RC* relationship equations derived from the QSSA equations (Fig. 2b (iii); see eq (4), (5)), to calculate the exact formula of *RC* with the weaker binding affinity. Taken together, we derived the exact formula for both drug-receptor complexes (*RC*_1_ and *RC*_2_), obtaining the exact formula of competitive qTMDD.

**Figure 2.**
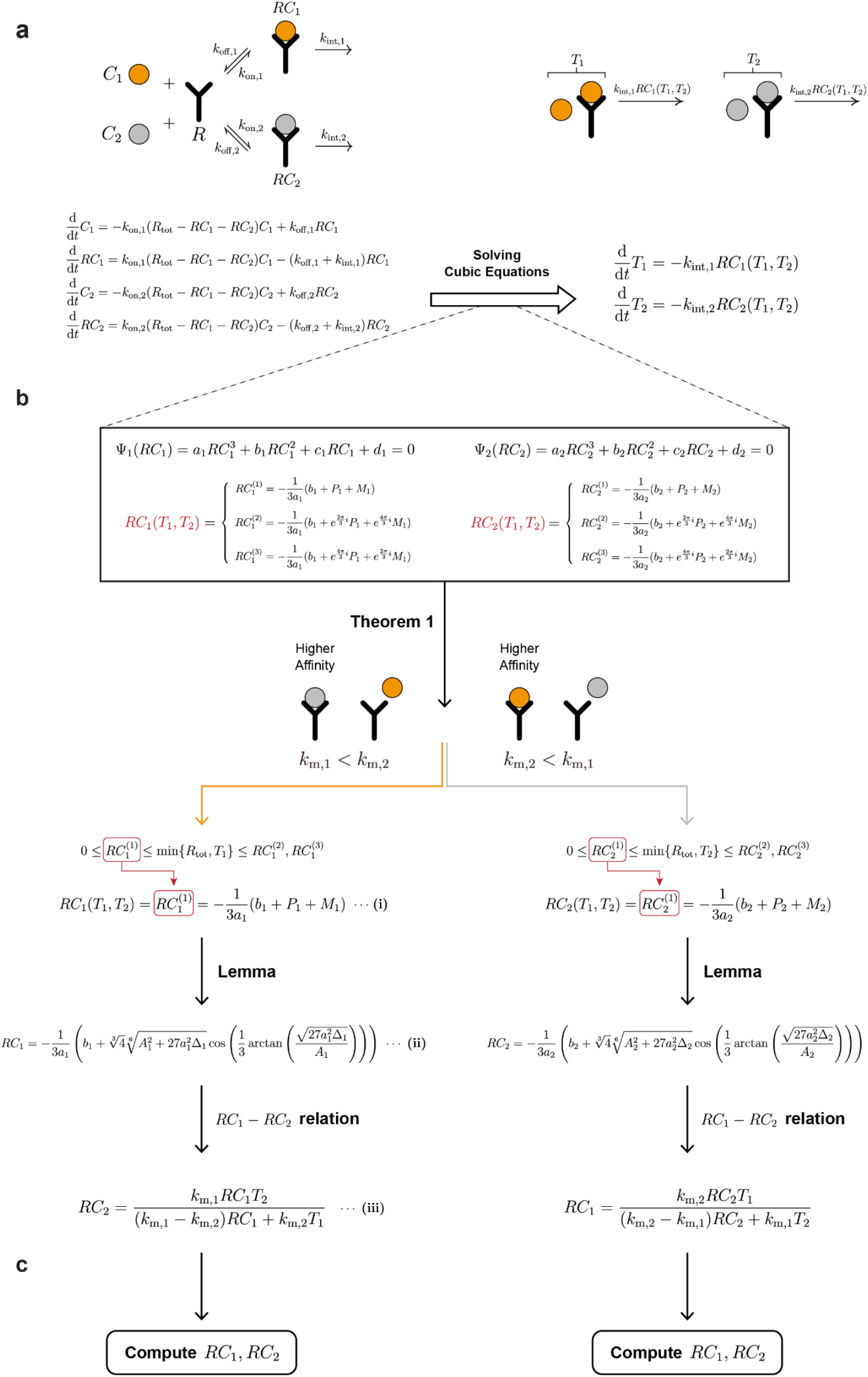
The biologically meaningful solutions for competitive qTMDD can be explicitly derived. **(a)** To simplify the full competitive TMDD model to the competitive qTMDD model, we provide a mathematical analysis for solving cubic equations and identify biologically meaningful solutions from the candidate solutions. **(b)** Deriving the competitive qTMDD model requires solving two symmetric cubic equations Ψ_1_(*RC*_1_) = 0 and Ψ_2_(*RC*_2_) = 0, which yield three candidate solutions each for *RC*_1_and *RC*_2_ (shown in the box). To identify the biologically meaningful solutions, we derive inequalities among three candidate solutions, as proven in Theorem 1 (see Methods for more details). Theorem 1 demonstrates that when the binding of *C*_1_(or *C*_2_) and *R* is stronger than that of *C*_2_ (or *C*_1_) and *R*, the unique biologically meaningful solution for *RC*_1_ (or *RC*_2_) is the smallest solution (i.e. 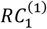 or 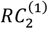) (i). This solution, initially expressed with complex terms, can be further simplified into real terms using the Lemma (ii) (see Methods for more details). Finally, the formula for*RC*_1_ (or *RC*_2_) directly provides the formula for *RC*_2_ (or *RC*_1_) through the *RC*_1_-*RC*_2_ relation (iii).

### The exact formula provides efficient computation of competitive qTMDD

We evaluated whether the exact formula enables faster computation of competitive qTMDD compared to previous methods–the bisection method and the comparison method. To do this, we measured the time required to compute *RC*_1_, *RC*_2_ (i.e., the roots of the cubic equation Fig. 1a (iv)) under various parameter conditions (Fig. 3a). In particular, 100 parameter combinations were generated by independently sampling each of the five different parameters (*k*_*m*,1_, *k*_*m*,2_, *T*_1_, *T*_2_, *R*_*tot*_) from the uniform distribution *U*(0, 1000). For each parameter combination, *RC*_1_ *and RC*_2_ were computed 50,000 times using each of the three methods: the exact formula, the bisection method, and the comparison method (see Methods for more details). This repetition was performed to ensure stable measurement of computation time, minimizing the effect of variability from single runs. The total computation times were visualized using violin plots (Fig. 3b).

**Figure 3.**
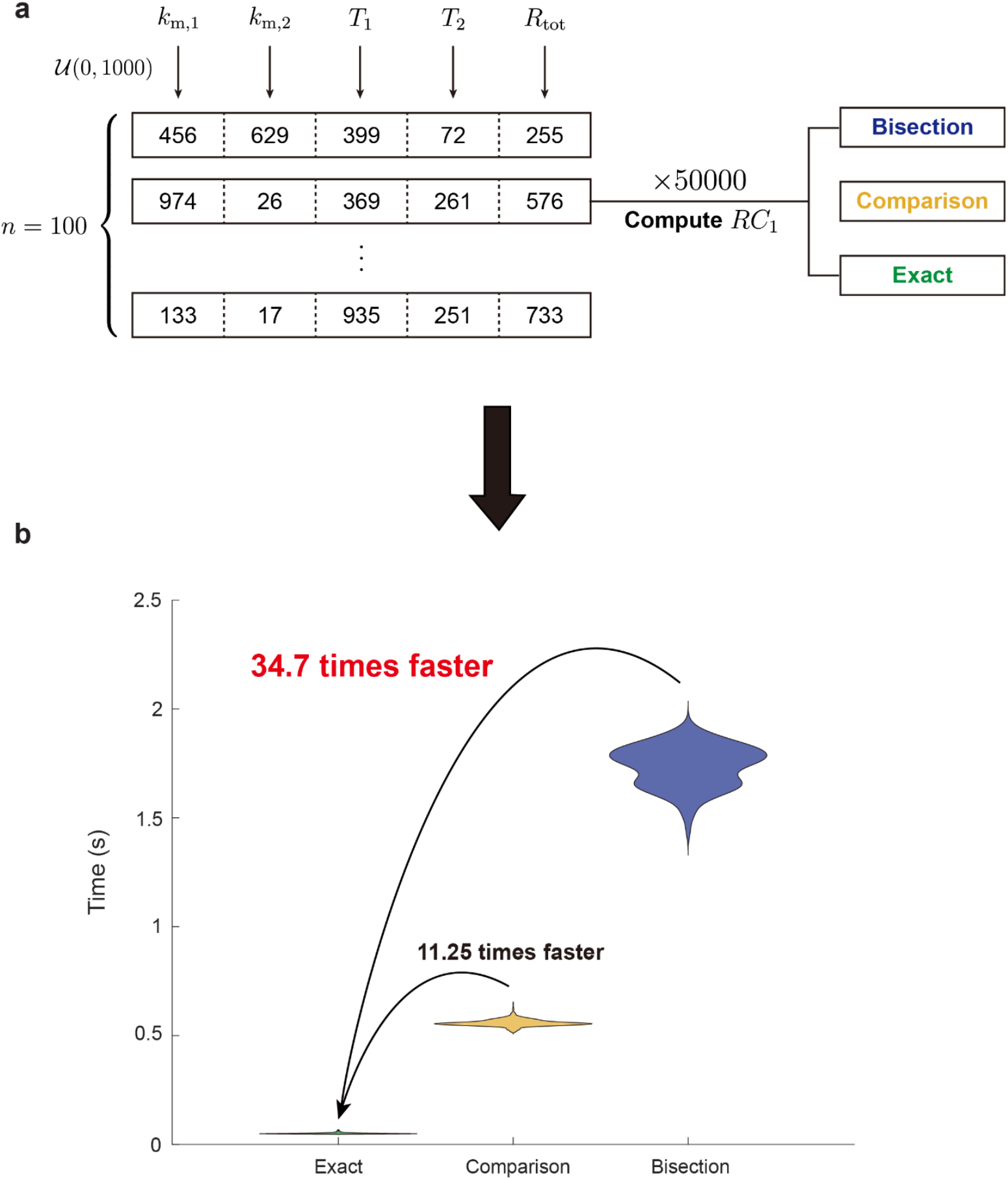
The exact formula provides the most efficient way of computing competitive qTMDD. **(a)** To compare the computation time in various parameter conditions, we generated 100 random parameter sets by randomly sampling each parameter from the uniform distribution: *U*(0, 1000). Then, we have obtained a distribution of computation time by measuring the time it takes to compute *RC*_1_ via tQSSA using each method (exact formula, comparison method, bisection method) under each parameter set. **(b)** Computation of competitive qTMDD using the exact formula is 11.25 times faster than the comparison method and 34.7 times faster than the bisection method.

The results show that the exact formula significantly improves computational efficiency (Fig. 3b). Specifically, it achieved a 34.7-fold speedup over the bisection method and an 11.25-fold speedup over the comparison method (see Methods for more details). Taken together, these results demonstrate that the exact formula significantly enhances the practicality of applying competitive qTMDD by drastically reducing computation time.

### The exact formula of competitive qTMDD allows efficient and unbiased Bayesian inference for competitive TMDD

The rapid computation of competitive qTMDD with the exact formula opens the door to its use in Bayesian inference, which was previously impractical due to the high cost of iterative computations. To demonstrate this potential, we used Bayesian inference to estimate the first Michaelis constant in competitive TMDD (*k*_*m*,1_) (Fig. 4a; see Methods for more details). We first generated the synthetic data of the full competitive TMDD model with a Gillespie algorithm (see Methods for more details). This synthetic data was then used to estimate *k*_*m*,1_ by fitting two reduced models: competitive mTMDD and qTMDD.

**Figure 4.**
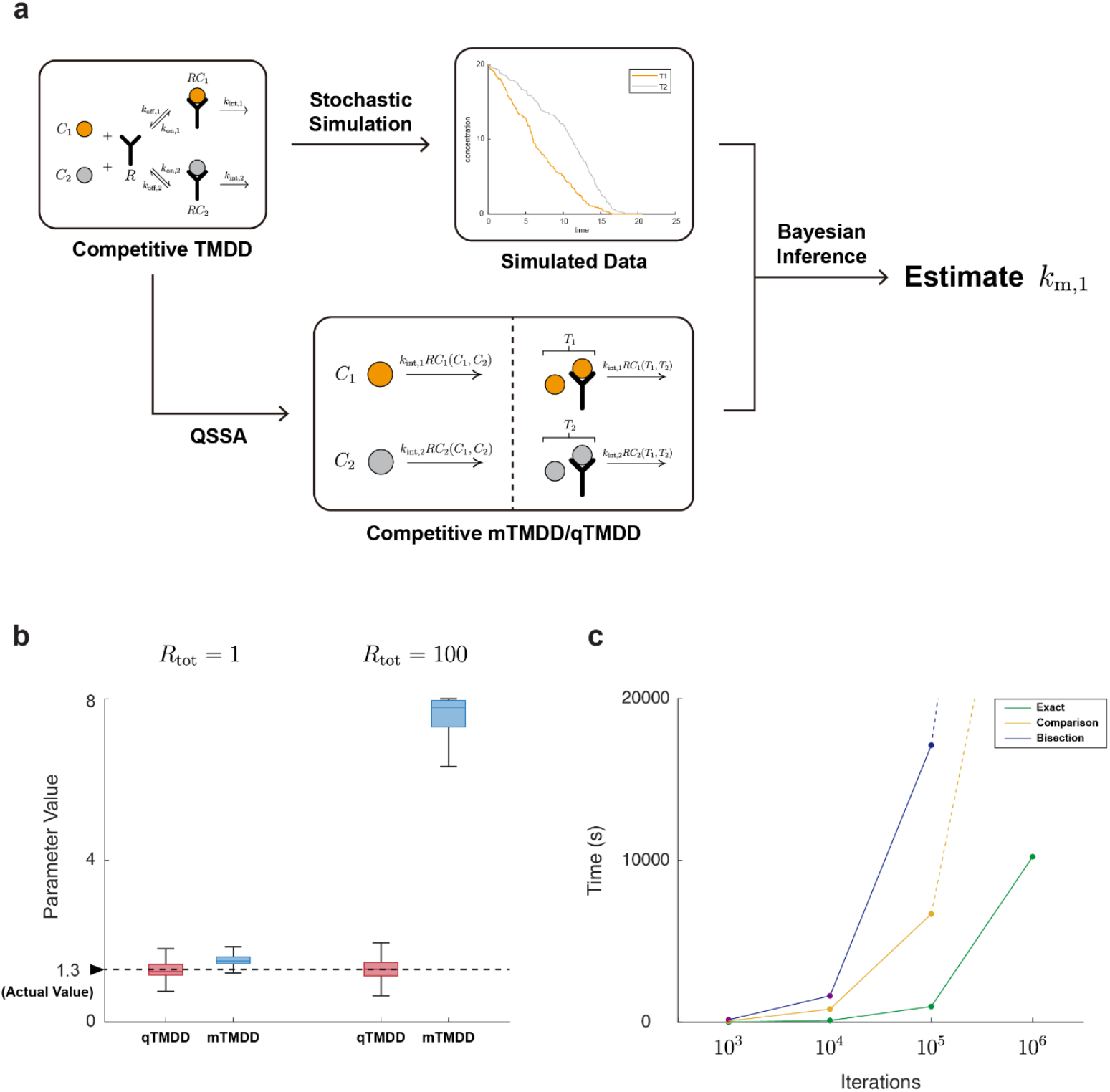
The exact formula provides an unbiased and efficient method of estimating parameters in a competitive qTMDD system. **(a)** We have obtained the simulated stochastic data using the Gillespie algorithm in a competitive TMDD system. To derive the likelihood of the simulated data, we have used both competitive mTMDD and qTMDD. Using the likelihood, we have estimated *k*_*m*,1_ via Bayesian inference. **(b)** Parameter estimation using competitive qTMDD (red) provides unbiased inference results under all conditions. Meanwhile, competitive qTMDD (blue) provides biased estimation results, especially when total receptor concentration (*R*_tot_) is high. **(c)** Use of the exact formula (green) is the most efficient way to estimate parameters in competitive qTMDD, rather than the comparison method (yellow) or bisection method (blue).

While *k*_*m*,1_ was accurately estimated using competitive qTMDD (Fig. 4b), inference based on the competitive mTMDD overestimated the value of *k*_*m*,1_ under high receptor concentration conditions (Fig. 4b, right). This indicates that the use of competitive qTMDD is crucial for accurate parameter inference. However, applying the competitive qTMDD with previous numerical methods requires a substantial computational cost (Fig. 4c). In particular, 1.8 hours and 5.6 hours were required when using the comparison method and the bisection method, respectively, to complete 10^5^ iterations of MCMC sampling. In contrast, the explicit formula reduced this time to approximately 0.25 hours. These results highlight that the explicit formula for competitive qTMDD is not only essential for accuracy but also enables practical use of Bayesian inference by significantly reducing computation time.

### The exact formula of competitive qTMDD estimates pharmacological parameters accurately and efficiently with real-world clinical data

Fast and accurate parameter estimation using the exact formula of the competitive qTMDD model shows its potential for estimating parameters in real-world clinical data. To investigate this potential, we estimated the pharmacological parameters by fitting full TMDD, mTMDD, and the exact formula of qTMDD to two representative real-world competitive TMDD cases, in which an endogenous and an exogenous substance compete for receptor binding: Case 1 for Anakinra and Case 2 for rhIL-7-hyFc. In Case 2, additional parameters for FcRn-mediated recycling and intramuscular dosing were included to account for the pharmacokinetic properties specific to rhIL-7-hyFc (see Methods for more details).

In Case 1, the ratio (*C*_0_ + *K*_*M*_)/*R*_tot_ for the endogenous drug is less than 1; thus, we expected that the mTMDD approximation would be invalid, whereas the qTMDD model would remain valid. Indeed, estimates of V_c_, k_a_, IIV_V_c,_ and IIV_k_a_ from the mTMDD model differed from those estimated by full TMDD, whereas qTMDD yielded parameter values close to those of the full TMDD model (Fig. 5a), despite comparable goodness-of-fit (GOF plots; S2 Fig) and predictive performance (VPC plots; S3 Fig). For example, the estimated V_c_ was 52.9 when using qTMDD and full TMDD, while mTMDD yielded 22.2 (S1 Table).

**Figure 5.**
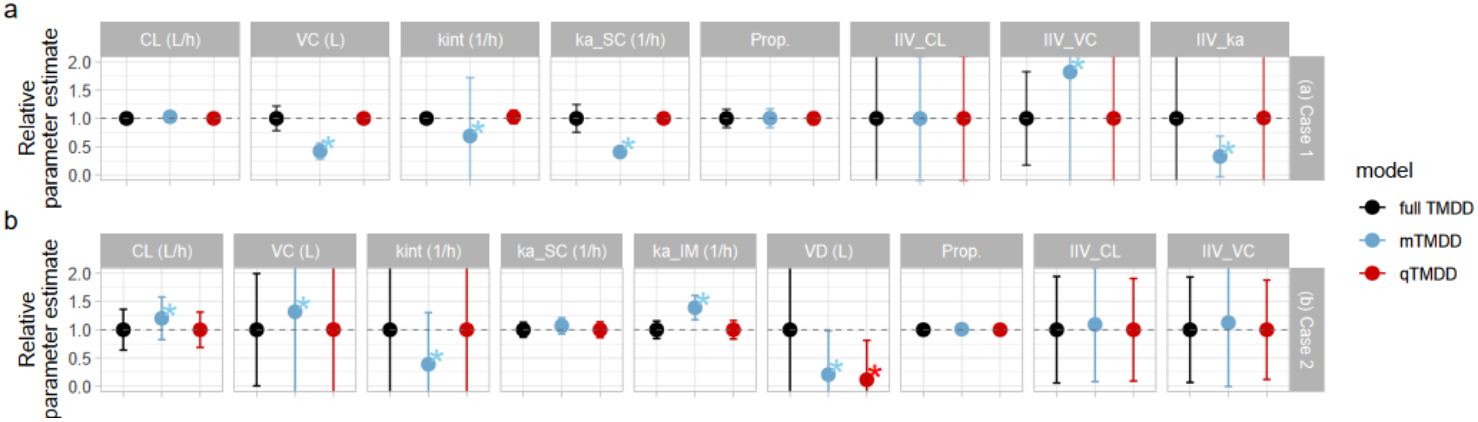
The competitive qTMDD yields parameter estimates similar to those of full competitive TMDD in real-world cases. **(a)** Plots of relative parameter estimates in Case 1 using full TMDD (black), mTMDD (blue), qTMDD (red). Full TMDD and qTMDD yield similar estimates for V_c_, k_int_, k_a_SC_, IIV_V_c,_ and IIV_k_a_. **(b)** Plots of relative parameter estimates in Case 2 using full TMDD (black), mTMDD (blue), and qTMDD (red). Full TMDD and qTMDD yield similar estimates for CL, V_c_, k_int,_ and k_a_IM_. CL: Apparent clearance of drug at central compartment, V_c_: Apparent distribution of drug and endogenous compound at central compound, k_a_: Absorption rate constant of drug from injection site (IM for intramuscular injection, SC for subcutaneous injection), k_int_: elimination rate of drug-receptor complex, Prop.: Proportional error, and IIV: Inter-individual variability. Asterisks (*) indicate parameters whose estimates in the approximation model deviated from those of the full TMDD model. Dots indicate relative parameter estimates, and error bars represent the corresponding 95% confidence intervals (CIs), calculated as CI = (Estimate/baseline) ± 1.96 * (SE / baseline). The dashed line at 1 indicates the full TMDD reference.

In Case 2, the ratio (*C*_0_ + *K*_*M*_)/*R*_tot_ is greater than 1 for both the endogenous and exogenous drugs. Thus, we expected that both the mTMDD and qTMDD models would be valid. However, unexpectedly, estimates of CL, V_c_, k_a_IM_, k_a_SC_, k_int_, IIV_CL, and IIV_V_c_ from the mTMDD model still deviated from those of full TMDD, while qTMDD yielded similar estimates (Fig. 5b). Furthermore, the objective function value (OFV) of the mTMDD model was lower than that of the qTMDD model (S1 Table), while GOF plots and VPCs were comparable.

To investigate the reason for this discrepancy between validity condition and parameter estimation results from the mTMDD model in Case 2, we generated a denser dataset by interpolating the original data using the fitted full TMDD model and compared the parameter estimates from the original sparse dataset (410 observations) and the interpolated dense dataset (1440 observations) (Fig. 6). With the denser dataset, parameter estimates from the mTMDD model became closely aligned with those from full TMDD model (Fig. 6a and S2 Table). In particular, the OFV value from mTMDD—initially low with the sparse data—increased progressively as more interpolated data were added (Fig. 6b). These results indicate that the mTMDD model was originally trapped in a false positive due to insufficient data sampling, and that increased sampling density mitigated this issue by reducing such false minima in the loss landscape, thereby enabling convergence toward the true solution. Taken together, these results show that—even when the theoretical validity conditions are satisfied—mTMDD can still yield inaccurate parameter estimates under limited sampling density. Therefore, qTMDD should be preferred when stable and accurate parameter estimation is required

**Figure 6.**
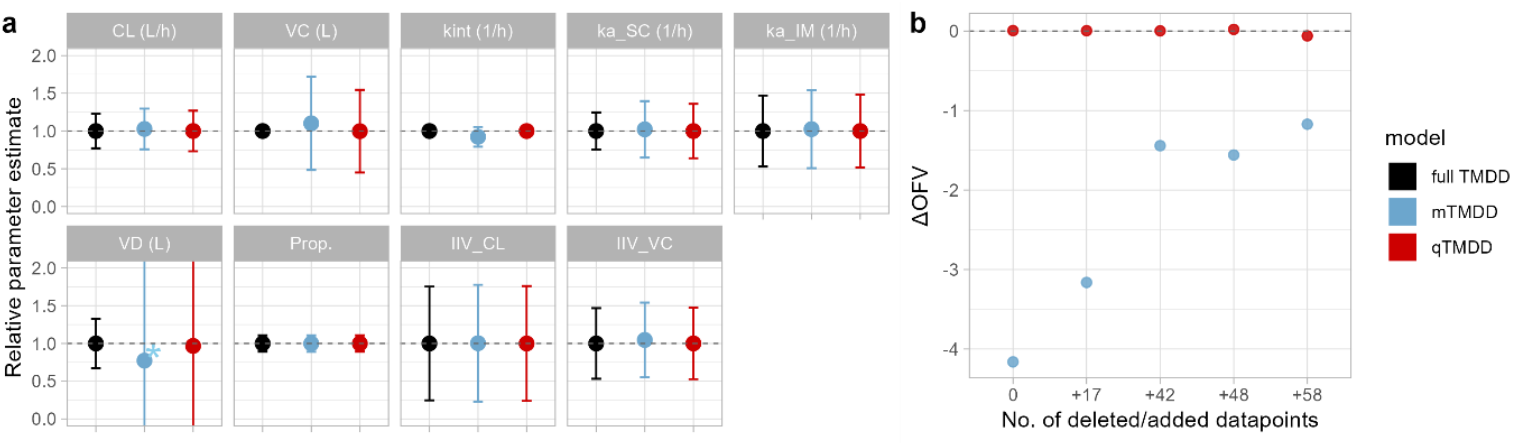
qTMDD yields parameter estimates similar to those of full competitive TMDD regardless of collected timepoint. **(a)** Plots of relative parameter estimates in densified data using full TMDD (black), mTMDD (blue), qTMDD (red). mTMDD showed decreased difference from full TMDD compared to the original data. Refer to Figure 5 for parameter definitions. **(b)** Effect of adding data points on the objective function value (OFV). For mTMDD, the OFV was 4.162 lower than that of the full TMDD at baseline; this difference narrows as the dataset is densified. ΔOFV is defined as OFV of the approximation model - OFV of the full TMDD model. See Methods for details of the dataset densification/reduction procedure.

Next, we investigated the computational advantage of qTMDD compared to the full TMDD model. To evaluate computational efficiency, we calculated the time required for estimating parameters over 10 repeated runs (estimation time) and the time required for estimating 1,000 resampled datasets with successful convergence to account for dataset-dependent variability (bootstrap time) (see Methods for more details). The results show that both estimation time and bootstrap time are shorter when using qTMDD than when using full TMDD (Fig. 7). In particular, in case 1, estimation time was reduced to approximately 28–92% of that required by the full TMDD model, and bootstrap time was reduced to about 30–43% (Fig. 7), despite requiring longer runtime than mTMDD (S1 Table). These results highlight that qTMDD offers a practical and more efficient alternative to full TMDD for real-world pharmacokinetic analyses.

**Figure 7.**
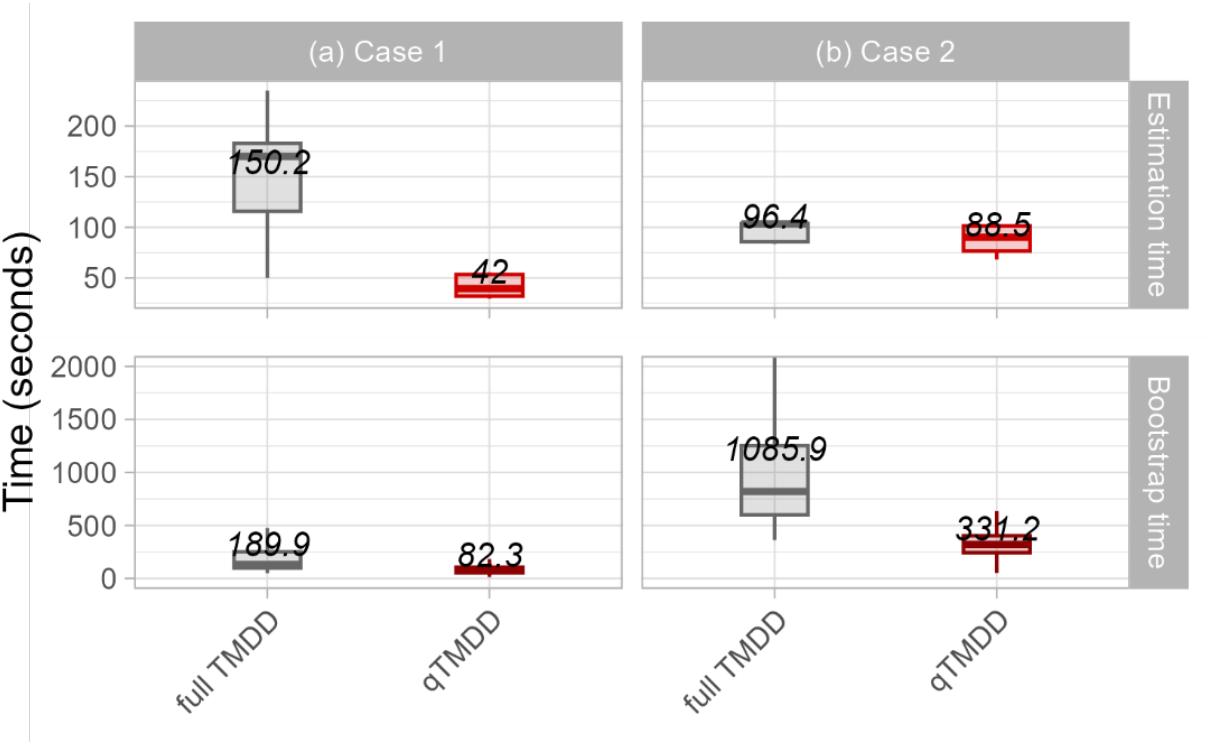
Competitive qTMDD enables faster parameter estimation than full competitive TMDD. Boxplots of estimation time (top) and bootstrap time (bottom) in Case 1 (a) and Case 2 (b) using full TMDD (grey) and qTMDD (green). Estimation times were obtained from 10 repeated runs of the final convergent model, while bootstrap times were obtained from 1,000 resampled datasets (successful runs only). For both estimation time and bootstrap time, qTMDD is faster than full TMDD. In the box plot, the line represents the median, the box represents the interquartile range, and the whiskers represent outliers.

## Discussion

Although competitive qTMDD is essential to accurately approximate competitive TMDD system, its derivation requires solving a cubic equation (Fig. 1c). While this can be done by using a cubic formula, the biologically meaningful solution among the three roots of the cubic equation was unknown, leading to the use of inefficient approximation methods such as the bisection or the comparison method (Fig. 1d). To address this problem, we derived the unique biologically meaningful solution, the exact formula of competitive qTMDD, by utilizing the analytic properties of cubic functions and the geometric properties of the complex plane (Fig. 2). This exact formula enabled competitive qTMDD simulations that were 11 times faster compared to previous approximation methods (Fig. 3). These rapid simulations enabled Bayesian inference using competitive qTMDD, and applying this to the simulation data resulted in more accurate and unbiased pharmacological parameter estimation than competitive mTMDD (Fig. 4). Furthermore, use of the exact formula of competitive qTMDD in real-world data enabled faster parameter estimation while yielding parameters similar to those estimated by the full TMDD model (Fig. 5). These results demonstrate that the exact formula of competitive qTMDD allows accurate and efficient pharmacological parameter estimation.

Parameter estimation using competitive qTMDD is likely to play an important role in pharmacometrics. With the rapid growth in biologics development, the use of TMDD models has expanded quickly. In clinical applications, it has often been assumed that exogenous drug concentrations are much higher than those of endogenous ligands, allowing competitive effects to be ignored. However, this assumption breaks down in several clinical scenarios. Examples include switching between soluble epoxide hydrolase inhibitors (31) and IVIG administered before and after infliximab (32) where effective concentration ranges partly overlap or prior exposure affects target saturation. In such cases, mechanism-based competitive TMDD is likely to better explain the observed nonlinearity, accumulation, and distributional changes, underscoring the need for a competitive TMDD model. Given this context, an exact formula for competitive qTMDD—one that accurately approximates the full competitive TMDD model while preserving computational efficiency—offers clear advantages for practical pharmacometric analyses. In particular, this accurate approximation includes robustness to limited sampling density: even when the theoretical validity conditions for mTMDD were satisfied, sparse sampling led mTMDD to biased parameter estimates and apparent local minima, whereas competitive qTMDD remained closely aligned with the full TMDD model in both sparse and densified datasets (Fig. 6). This robustness is important in clinical pharmacokinetic studies, where the number and timing of samples are constrained by ethical and logistical considerations. In terms of implementation, previous formulations of competitive qTMDD relied on complex-valued solutions or decision trees, which made it difficult to implement them in standard tools such as NONMEM. In contrast, our exact formula can easily be integrated into conventional PopPK/PD workflows. This enables direct estimation and simulation within population pharmacokinetic software, allowing model-based dose selection and scenario prediction to be immediately applicable in real clinical settings where competitive interactions are present.

Beyond the TMDD systems, the tQSSA-based reduction has also improved the accuracy of drug metabolism predictions, including applications in predicting hepatic intrinsic clearance and assessing drug-drug interactions mediated by the cytochrome P450 (CYP) induction (33–36). While these applications are grounded on a single enzyme kinetics where tQSSA derivation is straightforward, extending this framework to systems involving multiple enzymes results in a complex mathematical problem requiring the solution of a cubic equation—analogous to the competitive TMDD case addressed in this study (23, 37). We anticipate that our approach can be similarly applied to derive tQSSA reductions for multiple-enzyme drug metabolic systems, and investigating this potential with real-world data remains a promising avenue for future work.

Another important advantage of adopting the tQSSA-based reduction is that it resolves the structural limitations of sQSSA: satisfying the validity condition does not guarantee accurate parameter estimation, as shown in Fig. 5b. In particular, Choi et al. addressed this structural identifiability issue by leveraging the wide validity of tQSSA to integrate data across various experimental conditions and implement a sequential experimental design (38, 39). In this framework, parameter estimates derived from an initial dataset are utilized to determine the optimal initial concentrations for subsequent experiments, thereby maximizing the precision of the final estimation. Adopting this tQSSA-driven sequential strategy for both competitive TMDD systems and multiple enzyme drug metabolic systems would allow for a more accurate determination of kinetic parameters in complex competitive environments, representing a critical direction for future research.

Despite the advantages of competitive qTMDD, the high complexity of the exact formula makes its intuitive interpretation difficult (see Theorem 1 in Methods). Similar to the simplification of qTMDD to pTMDD in a non-competitive case (7), future work on simplifying the competitive qTMDD formulation will be important for broader adoption. Another potential direction of future work involves extending the framework to scenarios in which three or more drugs compete for a single receptor. In such situations, generalizing the derivation used for the two-drug case to a multi-drug competitive system will be necessary to obtain an exact competitive qTMDD formula. In addition, extending our approach to a stochastic and spatially distributed system represents a promising direction. Notably, stochastic systems necessitate different validity conditions for QSSA compared to deterministic ones (40–43), and spatial heterogeneity is known to alter the validity of the Michaelis-Menten rate law and QSSA in PDE models versus well-mixed ODE models (44, 45). Given this significant impact of stochasticity and spatial heterogeneity on QSSA, extending our framework to encompass these conditions would broaden its applicability to a wider range of realistic biological scenarios

There are also limitations when applying competitive qTMDD to real-world data in our analysis. In the dataset analyzed here, exogenous drugs are administered at much higher concentrations than endogenous substances, resulting in limited information about endogenous dynamics. Conducting similar analyses in settings where exogenous and endogenous concentrations are of comparable magnitude would help further validate the utility of competitive qTMDD across a broader range of clinically relevant scenarios.

## Methods

### Derivation of competitive mTMDD and qTMDD equations

Competitive TMDD is represented by the system of ODEs following (1, 7):

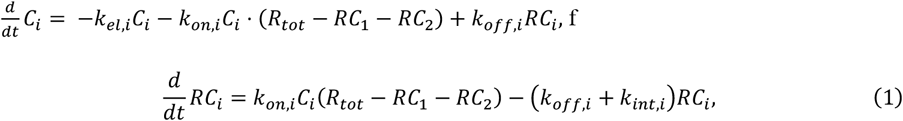

for *i* = 1,2, where *C*_*i*_ is the drug concentration, *RC*_*i*_ is the drug-receptor complex concentration, *R*_*tot*_ is the total receptor concentration, *k*_*el,i*_ is the drug elimination constant, *k*_*on,i*_ is the association rate constant, *k*_*off,i*_ is the dissociation rate constant, and *k*_*int,i*_ is the drug-receptor complex internalization constant.

To derive the competitive mTMDD, we assume that the drug-receptor complex concentration *RC*_*i*_ changes rapidly compared to the drug concentration *C*_*i*_ and thus reaches the QSS where *dRC*_*i*_/*dt* = 0.

Then, we obtain the following system of linear equations in *RC*_1_ and *RC*_2_:

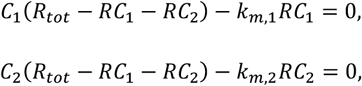

where *k*_*m,i*_ = (*k*_*off,i*_ + *k*_*int,i*_)/*k*_*on,i*_ for each *i* = 1,2. This system of linear equations can be solved easily as the following:

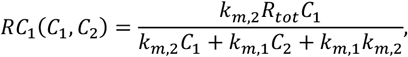

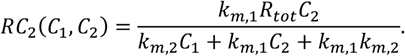

To derive the competitive qTMDD system, we assume that the drug-receptor complex concentration *RC*_*i*_ changes rapidly compared to the total drug concentration *T*_*i*_ = *C*_*i*_ + *RC*_*i*_ and thus reaches the QSS where *dRC*_*i*_/*dt* = 0 as done in the previous studies (1, 7). By substituting *C*_*i*_ with *T*_*i*_ − *RC*_*i*_ in the equation (1), we obtain the following system of quadratic equations in *RC*_1_ and *RC*_2_:

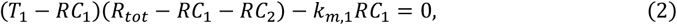

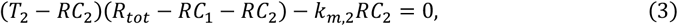

where *k*_*m,i*_ = (*k*_*off,i*_ + *k*_*int,i*_)/*k*_*on,i*_ for each *i* = 1,2. By simplifying equations (2) and (3), *RC*_1_ and *RC*_2_ can be expressed in terms of each other as follows (*RC*_1_-*RC*_2_ relation):

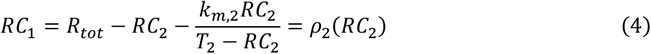

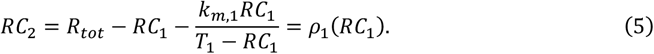

By substituting equations (4), (5) into equations (2), (3), we obtain two cubic equations Ψ_1_(*RC*_1_) = 0, Ψ_2_(*RC*_2_) = 0 where:

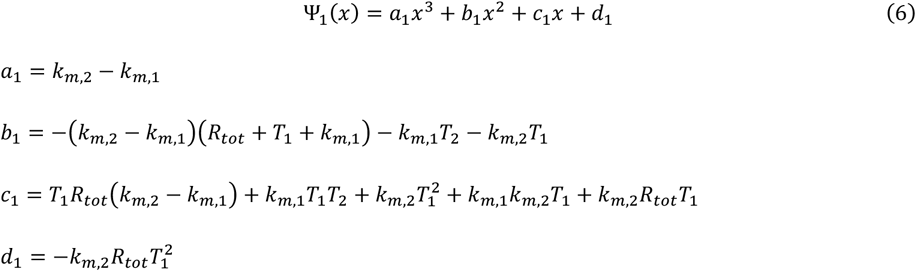

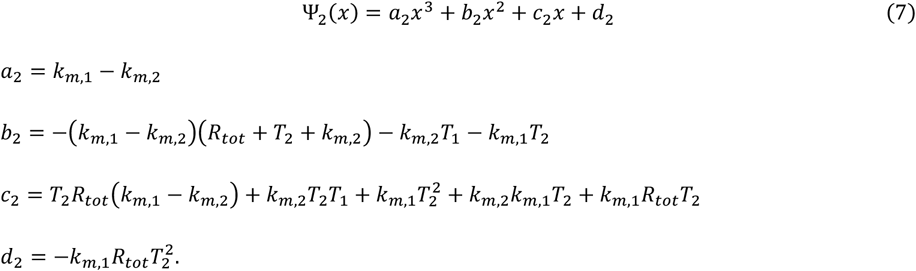

Solving this cubic equation and finding the biologically meaningful solution yields the exact formula of competitive qTMDD.

### Implementation of the bisection method

The bisection method computes the biologically meaningful root of the competitive qTMDD by approximating the root of Ψ_1_(*x*) in the biologically meaningful region ([0, min{*R*_*tot*_, *T*_1_}]) by reducing the interval through several iterations. The pseudo-code for the method can be found in Algorithm 1 in S1 Text.

Here, the error bound was set to ε = 10^−16^ and the maximum iteration *n* was set to 54, following a previous study (1). The MATLAB code for the bisection method can be found in S1 Text.

### Implementation of the comparison method

The comparison method computes all three roots of Ψ_1_(*x*) using the ‘roots’ function in MATLAB, and finds the biologically meaningful solution by checking whether each solution is in the biologically meaningful region [0, min{*R*_*tot*_,*T*_1_}]. While three roots of Ψ_1_(*x*) can be computed directly using the cubic formula, this method requires the use of software that supports complex number arithmetic since the cubic formula contains complex-valued terms. Therefore, we implemented the comparison method using the ‘roots’ function in MATLAB as done in (1). The MATLAB code for the comparison method can be found in S1 Text.

### Derivation of a biologically meaningful root for competitive qTMDD

To find a biologically meaningful solution of Ψ_1_(*RC*_1_) = 0, Ψ_2_(*RC*_2_) = 0, which is the solution satisfying (*RC*_1_, *RC*_2_) ∈ [0, min{*R*_*tot*_, *T*_1_}] × [0, min{*R*_*tot*_, *T*_2_}], we considered three different cases: *k*_*m*_,1 = *k*_*m*_,2_*m*_,1 < *k*_*m*_,2*, k*_*m*_,2 < *k*_*m*,1_.

If *k*_*m*,1_ = *k*_*m*,2_ = *k*_*m*_, then Ψ_1_(*x*) and Ψ_2_(*x*) become quadratic equations as *a*_1_ and *a*_2_ in (6) and (7) are zero. Therefore, the biologically meaningful solution of Ψ_1_(*x*) = 0 can be easily identified using the quadratic formula (see Case 1 of S1 Text for more details).

If *k*_*m*,1_ < *k*_*m*,2_, Yan et al. proved the existence of three distinct real roots of Ψ_1_ (*RC*_1_) = 0, and the smallest root is the unique biologically meaningful solution (1). (We simplified Yan et al.’s proof of these lemmas in Case 2 of S1 Text, using the signs of each coefficient of Ψ_1_(*x*), Ψ_2_(*x*)). However, finding the smallest root of Ψ_1_(*x*) was still challenging. To overcome this, we prove Theorem 1. The detailed proof can be found in Theorem 1 of S1 Text.

#### Theorem 1.

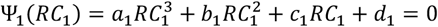 has a unique biologically meaningful solution

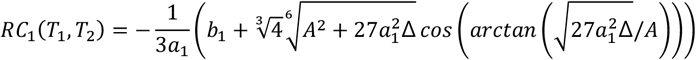

where 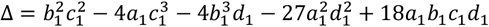 and 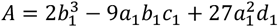.

Using both Theorem 1 and the relationship between *RC*_1_ and *RC*_2_ described in (4) and (5), the exact formula of *RC*_2_ can also be derived.

If *k*_*m*,1_ > *k*_*m*,2_ the same procedure can be applied by switching the roles of *RC*_1_ and *RC*_2_.

### Computational speed of simulating the competitive qTMDD model with different methods

To compare three computational approaches solving a competitive qTMDD model—the exact formula, the bisection method, and the comparison method—we evaluated their computational efficiency using randomly sampled parameter sets (Fig. 3). Specifically, 100 independent parameter sets (*k*_*m*,1_, *k*_*m*,2_, *T*_1_, *T*_2_, *R*_*tot*_) were sampled from *Uniform*(0,1000). For each parameter set, the total computation time required to calculate *RC*_1_, *RC*_2_ 50,000 times using each method was measured in MATLAB with the ‘tic-toc’ function to minimize the variability arising from single runs. The elapsed time was measured on a system with 8GB of RAM and an Apple M2 processor.

### Computational speed of Bayesian inference using competitive qTMDD with different methods

To compare the efficiency of three computational approaches solving competitive qTMDD with Bayesian inference, we have measured the computational time of Markov Chain Monte Carlo (MCMC) iterations (Fig. 4). First, the stochastic time series data for *T*_1_ and *T*_2_ were generated using the Gillespie algorithm for the full competitive TMDD system with parameters *k*_*el*,1_ = 0, *k*_*on*,1_ = 1, *k*_*off*,1_ = 0.3, *k*_*int*,1_ = 1 (i.e., *k*_*m*,2_ = 1.3) and *k*_*el*,2_ = 0, *k*_*on*,2_ = 0.3, *k*_*off*,2_ = 0.4, *k*_*int*,2_ = 2 (i.e., *k*_*m*,2_ = 8). Then we assumed that *k*_*el*,1_, *k*_*el*,2_, *k*_*m*,2_, *k*_*int*,1_, and *k*_*int*,2_ are known and estimated *k*_*m*,1_ using Bayesian inference (Fig. 4).

The likelihood of the observed data (i.e., the time series data for *T*_1_ and *T*_2_) was constructed via Poisson process. Suppose the time-series data of total drug concentration is given as 𝒟 = {( *t*_1_, *T*_1,1_, *T*_2,1_), (*t*_2_, *T*_1,2_, *T*_2,2_), ⋯, (*t*_*n*_, *T*_1,*n*_, *T*_2,*n*_)}. Since the rate of change of *T*_*i*_ is −*k*_*int,i*_*RC*_*i*_ (*i* = 1,2), for each *k* = 1,2, ⋯, *n* − 1, the decrement of *T*_*i*_ during [*t*_*k*_, *t*_*k*+1_] can be approximated by independent Poisson processes, each with rate *k*_*int,i*_*RC*_*i*_(*k*_*m*,1_, *k*_*m*,2_, *T*_1,*k*_, *T*_2,*k*_) ( *i* = 1,2), where *RC*_*i*_(*k*_*m*,1_, *k*_*m*,2_, *T*_1,*k*_, *T*_2,*k*_) is the *RC*_*i*_ value computed using competitive qTMDD.

From the previous assumption, for each *k* = 1,2, ⋯, *n* − 1 and *i* = 1,2:

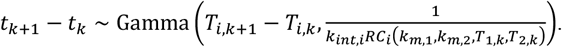

Therefore, the likelihood can be computed as:

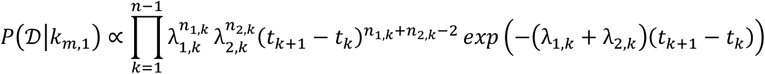

where λ_*i,k*_ = *k*_*int,i*_*RC*_*i*_(*k*_*m*,1_, *k*_*m*,2_, *T*_1,*k*_, *T*_2,*k*_), *n*_*i,k*_ = *T*_*i,k*+1_ − *T*_*i,k*_ (*i* = 1,2). By Bayes’ theorem, the posterior distribution is given as:

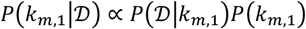

where *P*(*k*_*m*,1_) is the prior distribution of *k*_*m*,1_, *Uniform*(0, *k*_*m*,2_). The time for estimating *k*_*m*,1_ with 10^3^, 10^4^, 10^5^, 10^6^ MCMC iterations were measured for each of the three computational methods for solving qTMDD and their distributions were reported (Fig. 4). The elapsed time was measured on a system with 8GB of RAM and an M2 processor.

### Application of the exact formula of competitive qTMDD on real-world data

Two clinical cases were selected to evaluate approximation methods of competitive target-mediated drug disposition models. Case 1 involved competition between endogenous IL-1 and the exogenous drug anakinra at the IL-1 receptor, with 93 time–concentration observations fitted to the model. Case 2 involved competition between endogenous IL-7 and the exogenous drug rhIL-7-hyFc at the IL-7 receptor, with 410 time–concentration observations fitted. See (46) and (47) for detailed information. As described in the results section, C_0_ + K_M_ and R_tot_ should be compared, and we calculated (C_0_ + K_M_)/R_tot_ in each case for this purpose. In Case 1, this ratio was greater than 1 for the exogenous drug but less than 1 for the endogenous drug, suggesting that mTMDD would provide inaccurate approximations, whereas the exact formula of qTMDD would remain accurate. In Case 2, the ratio was greater than 1 for both the exogenous and the endogenous drugs, suggesting that both mTMDD and qTMDD would yield accurate approximations.

Previously established TMDD models describing drug–receptor interactions were adapted to incorporate competition between the exogenous drug and the endogenous ligand. Case 1 applied this framework to drug-receptor competition only (Fig.7(a)), whereas case 2 additionally included FcRn-mediated recycling to capture systemic disposition (Fig.7(b)). Full TMDD, mTMDD, and qTMDD formulations were applied to describe the binding of the drug and the endogenous ligand to the receptor, and their estimation accuracy and computational efficiency were compared. A population pharmacokinetic model was constructed by accounting for inter-individual and residual variability, and parameter optimization was performed using NONMEM 7.5 and PsN 5.3.1. To avoid over-parameterization, certain parameters were fixed to in vitro values reported previously: in case 1, k_el_endo_, K_SS_exo_, R_tot_, k_on_, k_on_endo_, and k_off_endo_; in case 2, k_on_exo_, k_off_exo_, K_SS_exo_, K_SS_FcRn_, R_tot_, FcRn, k_uptake_, and k_recycle_. To minimize the influence of the initial values on the parameter estimation, we used an option that varied the initial estimates over a 90% range around the final model, and most parameter estimates differed by less than 2% in both cases. For both cases, receptor concentration was assumed constant, with receptor degradation set equal to k_int_exo_ = k_int_endo_. In case 2, the endogenous ligand concentration was also assumed constant, with k_deg_endo_ = set equal to the same value.

**Figure 7.**
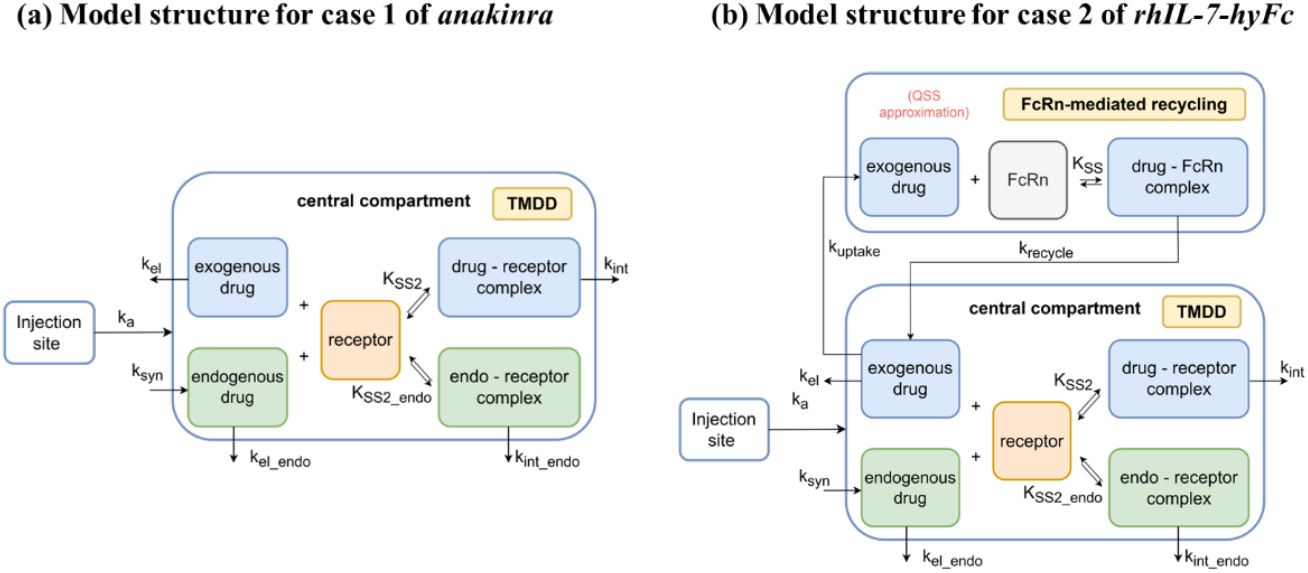
Model structure for cases 1 and 2. k_a_: absorption rate constant of drug from injection site, K_SS2_: equilibrium constant of drug and receptor binding, K_SS2_endo_: ~of drug and endogenous compound binding, K_SS1_: ~of drug and FcRn binding, CL: apparent clearance of drug at central compartment, V_c_: apparent distribution of drug and endogenous compound at central compound, k_int_: elimination rate of drug-receptor complex, k_syn_: synthesis rate of endogenous compound, k_el_endo_: elimination rate of unbound endogenous compound, k_int_endo_: elimination rate of drug-endogenous compound complex, V_D_: apparent distribution of drug at distribution site, k_uptake_: uptake rate constant of drug from central to distribution space, k_recycle_: recycling rate constant of drug from distribution space to central due to its binding with FcRn, receptor: therapeutic receptor of anakinra or rhIL-7-hyFc, FcRn: neonatal Fc receptor.

An additional analysis was conducted to assess whether the estimation results for case 2 depended on the data-collection time point. We defined a dense grid by interpolating the early phase at 0.1-h spacing from 0 to 2 h and 0.5-h spacing from 2 to 25 h, simulated concentration–time profiles using the final full TMDD model, and then augmented the original dataset by adding only simulated observations at times not present in the original data; when a time overlapped, the original observation was retained. This procedure produced the densest set (+58). Starting from these +58 datasets, we obtained less-dense variants by sequentially removing early timepoints: +48 with 0.2-h spacing from 0 to 2 h, +42 with 0.5-h spacing from 0 to 2 h, and +17 with 1-h spacing from 0 to 25 h. Each of these datasets was analyzed with the final full TMDD, mTMDD, and qTMDD models, and the corresponding objective function values (OFV) were compared to assess the impact of sampling density.

Performance comparisons were conducted to evaluate estimation accuracy and computational efficiency. Estimation accuracy was defined as the degree of agreement with the parameter estimates from the full TMDD model, which was regarded as the reference standard. For this, we compared parameter estimates, objective function values (OFV), residual-based diagnostic indices such as goodness-of-fit (GOF) plots, and simulation-based diagnostics such as visual predictive check (VPC) plots between approximation models and the full TMDD model. Computational efficiency was evaluated based on estimation and bootstrap times. Estimation time was measured by running the parameter estimation 10 times, with the initial condition set as the final parameter estimates shown in the S1 Table. Then, its distribution was reported to account for variability due to random fluctuations. For bootstrap time, we resampled the original data 1,000 times, and for each resample, estimated parameters starting from the values in S1 Table, reporting the distribution of the 1,000 estimation times. The elapsed time was measured on a system with 8 GB of RAM and an AMD Ryzen 7 5800H (3.2 GHz) processor for estimation, and on one with 256 GB of RAM and an AMD Ryzen Threadripper PRO 5965WX 24-core processor for bootstrap.

## Data Availability

The real-world data for Anakinra and rhIL-7-hyFc are presented in S3 Table and S4 Table, respectively.

## Code Availability

The simulation codes underlying this work are provided as a ZIP file ‘qTMDD.zip’, S1 File, for the reviewers and will be made publicly available upon acceptance. The NONMEM codes are available in S2 Text.

## Acknowledgements

The research was supported by the following agencies and institutions: the Institute for Basic Science (IBS-R029-C3, J.K.K.), Basic Science Research Program through the National Research Foundation of Korea(NRF) funded by the Ministry of Education (RS-2025-25397599, J.K.K., H.Y.), and the National Research Foundation of Korea(NRF) grant funded by the Korea government (MSIT; No. RS-2022-NR069643, H.Y.)

## Author Contributions Statement

Conceptualization: Dongju Lim, Hwi-yeol Yun, Jae Kyoung Kim

Data curation: Hwi-yeol Yun

Formal analysis: Jaeyun Cha, Taehong Kim, Dongju Lim, Hyeseon Jeon

Funding acquisition: Hwi-yeol Yun, Jae Kyoung Kim

Investigation: All authors

Methodology: Jaeyun Cha, Taehong Kim, Dongju Lim, Jae Kyoung Kim

Project administration: Dongju Lim, Hwi-yeol Yun, Jae Kyoung Kim

Supervision: Jae Kyoung Kim

Validation: All authors

Visualization: Jaeyun Cha, Taehong Kim, Hyeseon Jeon

Writing – original draft: All authors

Writing – review & editing: All authors

## Supporting information

**S1 Text. Supplementary materials for competitive qTMDD**

**S2 Text. The NONMEM code of the population pharmacokinetic model with each approximation method for competitive TMDD implemented**

**S1 Fig. Competitive qTMDD provides an accurate approximation of full competitive TMDD for all total receptor concentrations and initial drug concentrations**. (a) Each column represents total receptor concentration *R*_*tot*_ = 2, 20, 40 and each row represents initial drug concentration *C*_0_ = *C*_1,0_ = *C*_2,0_ = 2, 20, 40. Each curve represents the simulated total drug 1 concentration *T*_1_ = *C*_1_ + *RC*_1_ over time for *k*_*on*,1_ = 1, *k*_*off*,1_ = 0.5, *k*_*int*,1_ = 1.1, *k*_*el*,1_ = 0.01, *k*_*on*,2_ = 0.3, *k*_*off*,2_ = 0.4, *k*_*int*,2_ = 2, *k*_*el*,2_ = 0.01. (b) Competitive qTMDD (left, red dashed line) shows an accurate approximation to the full competitive TMDD (left, black solid line) for all total receptor concentrations and initial drug concentrations (*C*_1,0_ = *C*_2,0_ = *C*_0_). (c) In contrast, competitive mTMDD (right, blue dashed line) only shows a fairly good approximation when total receptor concentration is sufficiently smaller than the initial drug concentration.

**S2 Fig. Goodness-of-fit plots of (a) Case 1 and (b) Case 2**. (A, E) population prediction vs observations, (B, F) individual predictions vs observations, (C, G) time vs conditional weighted residuals, (D, H) population prediction vs conditional weighted residuals. The fitted lines of the observation vs. prediction closely matched the y=x line (A, B, E, F), and most conditional weighted residuals (CWRES) were within the range of −2 to 2 with the fitted line positioned close to zero (C, D, G, H).

**S3 Fig. Visual predictive checks of (a) Case 1 and (b) Case 2**. Observed 5^th^, 50^th^, 95^th^ percentiles are compared with 95^th^ prediction intervals from 200 simulations. No prediction correction was applied. This VPC plot showed that the prediction intervals adequately encompassed all observations without excessive overlap between intervals, indicating that the model appropriately captured both the central tendency and variability of the data.

**S1 Table. Parameter estimate result, OFV values and elapsed time**

**S2 Table. Parameter estimate results and OFV values for Case 2 estimation results with dense simulated dataset**

**S3 Table. Dataset for Case 1**

**S4 Table. Dataset for Case 2**

**S1 File. qTMDD.zip**

